# ProjectSVR: Mapping single-cell RNA-seq data to reference atlases by supported vector regression

**DOI:** 10.1101/2023.07.31.551202

**Authors:** Jianing Gao, Shipeng Guo, Yuanwei Zhang

## Abstract

The rapid accumulation of public single-cell RNA-seq (scRNA-seq) data and advances in integration algorithms make it possible to build the organ and body-scale cell atlases after overcoming the batch effect originating from different studies, donors, and sequence platforms. However, projecting the query cells onto the carefully constructed reference atlas to rapidly interpret cell states remains challenging. Here, we present ProjectSVR (https://github.com/jarninggau/ProjectSVR), a machine learning-based algorithm for mapping the query cells onto well-constructed reference embeddings. We demonstrate the effectiveness of ProjectSVR in multiple case studies of different tissue and experimental design, including (1) interpreting the decidual immune microenvironment from recurrent pregnancy loss (RPL) patients, (2) resolving the tumor-infiltrated T cell states from patients receiving anti-PD1 treatment, (3) mapping the genetic-perturbed mouse germ cells to integrated mouse testicular cell atlas (mTCA) for inferring their arrested developmental stage, and (4) evaluating in vitro induced meiosis by via mapping the induced germ cells onto mTCA. Taken together, our results show that ProjectSVR is a useful tool for reference mapping and make the scRNA-seq data analysis more convenient when well-annotated atlases emerge.

## Introduction

Advances in single-cell RNA sequencing (scRNA-seq) revolutionized the exploration of cellular heterogeneity during development and disease ^1^. With the advent of feasible scRNA-seq technology and decreasing sequencing costs, an increasing number of researchers have adopted this technique in their studies. However, the rapid accumulation of scRNA-seq data presents new challenges in achieving fast and reproducible biological interpretation. On one hand, typical scRNA-seq analysis pipeline follows an unsupervised approach, which includes normalization, feature selection, dimension reduction, clustering, and manual cell-type annotation, primarily due to the lack of appropriate reference atlases. These processes, especially manual cluster annotation, are subjective and time-consuming, thereby limiting the reproducibility of results. On the other hand, the development of data integration algorithms has led to the creation of comprehensive body- and organ-scale atlases ^2^, which offer an opportunity for consensus cell type annotation by incorporating their biological context ^3–5^. Nevertheless, mapping novel datasets onto well-constructed references still poses a significant challenging ^6^.

Recently, several supervised methods have been developed to address the aforementioned issues. These methods are primarily based on two main strategies: classification and projection ^6^. While the classification-based strategy has already demonstrated good performance in discrete cell type prediction and reached a certain level of maturity ^7–9^, it still has some limitations. These limitations include: (1) classification models cannot be used for dealing with continuous cell states; (2) models must be retrained when the reference cell labels are changed; and (3) it is challenging to balance the model performance and granularity of predicted labels, considering the recently published atlases using a multi-layer annotation system ^10^.

Projection-based strategy, also known as reference mapping, is a well-recognized method to remedy the limitations of classification methods. It involves placing query cells onto a well-constructed reference without corrupting the reference embeddings ^6^. Recently, several algorithms have been introduced to seamlessly integrate the steps of reference building and query mapping. Notable examples of such algorithms include Seurat ^11^, Symphony ^12^, scArches and ProjectTILs ^13^. These methods efficiently map query cells to reference embeddings by utilizing the compressed features, using fixed parameters generated during the reference-building step. However, data integration presents substantial computational difficulties, and the presence of batch structures can adversely affect the performance of integration methods^2, 14^. The joint reference building and query mapping process has hindered the application of cell atlases constructed using self-defined integration pipelines, exampled by the recently published pan-cancer tumor-infiltrating T cell landscape ^15^.

To address the above-mentioned challenge, we developed ProjectSVR, an R package for reference mapping based on supported vector regression (SVR). When a reference atlas is provided, ProjectSVR follows a two-step process for reference mapping: (1) Fitting a collection of SVR model ensembles to learn embeddings from feature scores of the reference atlas; (2) Projecting the query cells onto the consistent embeddings of the reference via trained SVR models. ProjectSVR can overcome the batch effect from different query datasets and assigns an identity to the consensus cells. The accuracy and effectiveness of ProjectSVR in mapping cells into multiple tissues have been extensively demonstrated through various case studies. In addition, we offer user-friendly interfaces and numerous pre-trained models to facilitate the interpretation of biological implications in scRNA-seq data across a wide range of biological fields.

## Results

### Directly predicting the reference embeddings from the raw count matrix using support vector regression

Reference mapping means localizing the query cells within integrated reference embeddings without requiring access to the reference data ^12^. Previous studies have employed modified data integration methods by fixing underlying parameters to construct a transform model that projects query data onto reference embedding space ^12, 13^. However, this approach is dependent on a specific data integration algorithm. As the reference embeddings describe the cell states, we investigated the feasibility of training a regression model from the raw count matrix to predict reference embeddings. Building upon this original idea of ProjectSVR, we initially extracted marker genes from a well-annotated reference atlas using canonical methods or consensus non-negative matrix factorization (cNMF) ^16^. Subsequently, these gene sets were utilized to calculate signature scores using the UCell algorithm ^17^ (Fig. 1A). This step compresses the raw count matrix, containing thousands of genes, into a signature matrix with fewer than one hundred dimensions. We then trained an SVR model using the signature matrix to predict reference embeddings. Considering the computational cost of SVR on large samples, we employed a subsampling strategy to train multiple models for each dimension of embeddings. The final predicted embeddings are obtained by taking the median of predictions make by all models (Fig. 1B). Once the integrated SVR model is trained, query data can be projected onto the reference by predicting the embeddings, and all the cell metadata from the reference will be transfered to the query data through neighborhood-based predictors (Fig. 1C).

**Figure 1.**
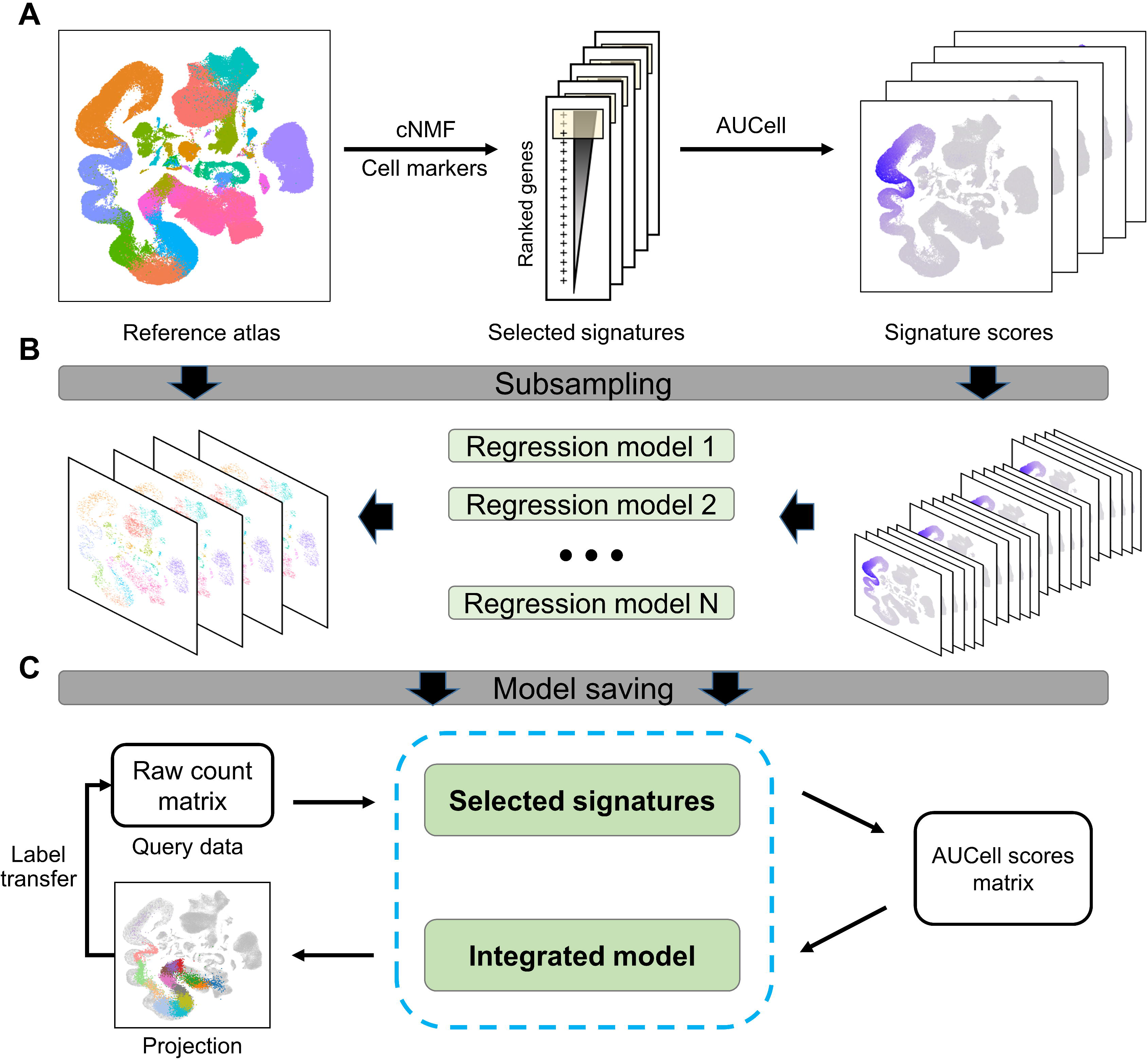
The ProjectSVR workflow. (A) The ProjectSVR takes integrated embeddings (e.g. UMAP) and gene set scores calculated via the UCell algorithm as inputs. The gene set is prepared by cell type-specific markers or high-weighted genes identified using consensus non-negative matrix factorization. (B) Then a regression model is fitted by supported vector regression (SVR). Because a reference atlas contains a hundred thousand cells but SVR is a small sample model, we subsampled cells into thousands of cells and trained a set of SVR models in parallel. (C) The selected gene set and models are saved for reference mapping. The query data is first transformed into a signature score matrix from the raw count matrix to project query data onto a reference atlas. Then the trained model is used to predict embeddings. ProjectSVR adopts the median value of the predictions as final outputs. Once the projection is finished, the cell labels from the reference are easily obtained by query cells using a k-nearest neighbors (KNN) classifier.

### Mapping the PBMC data across technologies to the harmonized reference space

We designed a PBMC task to assess the feasibility of ProjectSVR for reference mapping. A pre-build PBMC reference atlas is required for ProjectSVR. Therefore, we utilized the scVI ^18^ algorithm to integrate well-annotated blood samples from various studies collected in the DISCO database ^19^ and visualized cells by UMAP using scVI-generated latent space (Fig. 2A). Subsequently, we mapped 20,571 PBMCs obtained from three different 10x technologies (3’v1, 3’v2, and 5’) to this reference atlas, comparing the mapping results with *de novo* integration methods, both with and without batch correction. In order to evaluate the dimensionality reduction outcomes, we employed UMAP visualization and local inverse Simpson’s Index (LISI) ^20^. LISI quantifies the effective number of distinct categories represented in the local neighborhood of each cell based on category labels. When cells are well-mixed, the LISI scores will approach the actual number of category labels (in this case, three). Without batch effect correction, the UMAP embeddings of query PBMCs exhibited skewness primarily due to technological discrepancies rather than cell types (Fig. 2B and C, mean LISI = 1.0), indicating a considerable batch effect in the query data. However, by utilizing ProjectSVR, cells were projected into biologically coherent reference groups whilte removing technology-specific variations (mean LISI = 2.4). This achievement was comparable to *de novo* integration using harmony (mean LISI = 2.5, Fig. 2B-D). Furthermore, predicted embeddings based on ProjectSVR successfully separated pre-labeled cell subtypes, indicating biological meaningful results (Fig. 2D).

**Figure 2.**
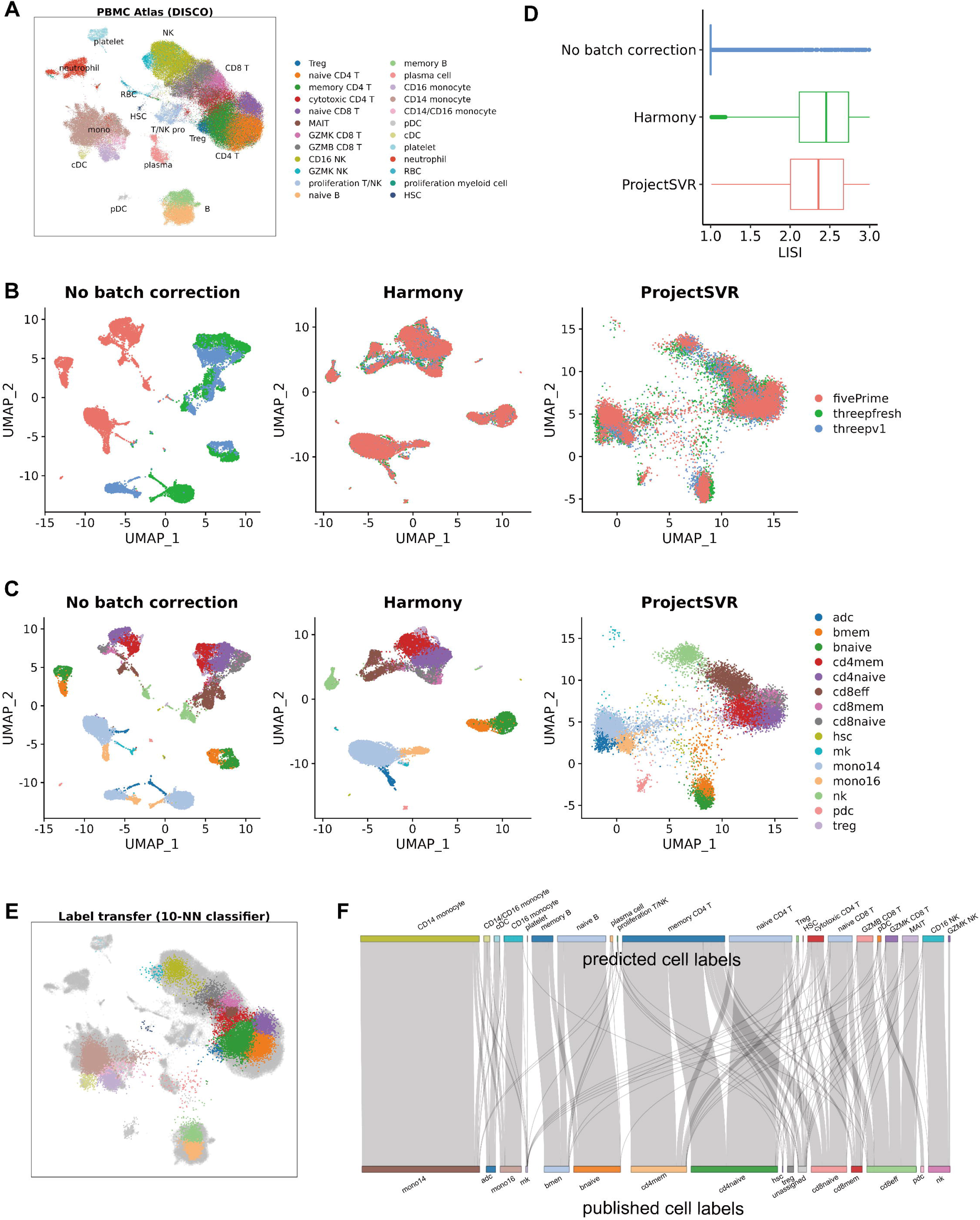
Mapping PBMC query data from different technologies to DISCO blood atlas. (A) Uniform manifold approximation and projection (UMAP) plots showing the blood atlas in DISCO database. Each dot represents a single cell, colored by cell type. (B-C) Projection of PBMC datasets from three 10x protocols onto DISCO blood atlas in (A) Left, UMAP plots without batch correction; middle, UMAP plots by harmony; right, query cells projected by ProjectSVR. Dots are colored by protocols (B) and published cell types identified by Korsunsky et al. (D) Boxplot showing the LISI (Local Inverse Simpson’s Index) distribution of data presented in (B). (E) Transfer cell labels from blood atlas (reference) to query cells using a 10-NN classifier on the ProjectSVR embedding. Query cells are colored by cell type defined in (A), and grey dots represent reference cells. (F) Sankey diagram shows the consistency between predicted cell labels in (E) and published cell labels in (C).

It is essential to develop metrics for evaluating the mapping quality, in order to assist users in identifying and discarding erroneous projections of query cells. A successful projection should preserve the local topological relationships of each query cell between the feature space and projection space. Therefore, we introduced a metric called mean k-nearest-neighbor distance (mean k-NN dist) to assess the mapping quality (Fig. S1A-C). First, a k-NN graph is constructed in the feature space (Fig. S1A). Then we calculate the average distance between each query cell and its k nearest neighbors using the projected embeddings. If a query cell is correctly projected, it will exhibit a similar local structure in both the feature and projection spaces, resulting in a small mean k-NN dist (Fig. S1B). Conversely, if the query cell is far from its nearest neighbors (as defined in the feature space) in the projection space, it will have a significantly larger mean k-NN dist (Fig. S1C). We calculated the mean k-NN dist for the query PBMCs and observed that only a small proportion of cells (2.6%) with a mean k-NN dist larger than 1.5 were classified as low-quality projections (Fig. S1D-F). These cells were dispersed away from cluster centers and subsequently excluded during label transferring (Fig. S1F).

A valid projection enables accurate cell type prediction. In this study, we utilized a simple k-NN classifier (k=10) to transfer annotated cell labels from the DISCO database to query cells. We then compared the predicted labels with the original report by Korsunsky et al. ^20^ (Fig. 2E). To evaluate the performance of cell type prediction, we harmonized the query and reference labels (Table S2). The k-NN classifier achieved high specificity (>0.95) for most cell subtypes, such as CD16 NK, pDC, and naïve B (Fig. 2F and Fig. S2A). However, the platelet and Treg clusters exhibited low sensitivity (0.20 and 0.21 respectively, Fig. S2A). To investigate further, we examined megakaryocytes (pre-labeled as ‘mk’) and found that only 16.7% of them were correctly predicted as ‘platelet’. The majority were ‘incrrectly’ classified as CD14 monocyte and T cells (53.5% and 12.5% respectively) (Fig. S2B). This suggests a potential contamination of these wrongly classifed cells. To verify this, we analyzed the expression of PPBP (a marker for platelets), CD14 (a marker for CD14 monocytes), and CD3D (a marker for T cells). The correctly classified platelets exhibited the highest levels of PPBP, while the ‘wrongly’ classified CD14 monocytes and T cells expressed their corresponding markers (Fig S2C). A similar scenario was observed for regulated T cells (Tregs) (Fig. S2D, E). These findings demonstrate that the projection results of ProjectSVR enable the identification of purer rare cell clusters, potentially due to the inclusion of a biological coherent reference and the cell-to-cell independence during the projection process. Additionally, we observed that the majority (81.4%) of effector CD8+ T cells (cd8eff) could be further classified into three subtypes based on correlated markers (Fig. S2F, G). This implies that ProjectSVR predicts biological meaningful embeddings and enables the discrimination of subtle cell subtypes.

### ProjectSVR facilitated the interpretation of the decidual immune microenvironment in RPL patients

It is a common scenario in scRNA-seq experiments to analyze paired samples of health and diseased individuals in order to explore changes in cell population distribution and perturbed trainscriptional programs. A successful reference mapping method should be capable of accurately assigning query cells to the reference, even when there are transcriptional perturbation in disease samples. In our study, we utilized ProjetSVR to project decidual immune cells from both healthy individuals and patients with recurrent pregnancy loss (RPL) ^21^ onto a previously reported maternal-fetal interface atlas to evaluate its efficacy ^4^. First, we integrated the reference using the fastMNN algorithm to correct for batch effects arising from different single-cell library construction protocols (Fig. 3A). Then, we employed ProjectSVR to learn an ensembled SVR model and projected query cells onto this reference. ProjectSVR effectively corrected the batch effects originating from different samples (Fig. S3A-C). Furthermore, the Louvain clustering algorithm was used to identify the types of query cells, which served as the true labels, and these cells were correctly placed into the corresponding cell groups in the reference atlas (Fig. 3B and Fig. S3C). We utilized a KNN classifier with ten nearest neighbors to transfer reference labels to query cells. The resulting confusion matrix demonstrated coherence between the true labels and the predicted labels (Fig. 3C). Notably, we observed that cluster 8 in the query data exhibited high expression of a set of genes encoding heat shock proteins (HSP) (Fig. S3D). However, the upregulation of HSP genes was attributed to a stress response during tissue dissociation and storage, rather than being an actual biological signal ^22^. We found that ProjectSVR effectively mitigated the transcriptome bias caused by this stress response, thereby enabling accurate prediction of the cell identity (Fig. 3D and Fig.S3E, F). Finally, we investigated the changes in the decidual natural killer (dNK) cell population between healthy controls and RPL patients. As reported in the previous study ^21^, our findings revealed a decrease in the dNK1 population and an increase in the dNK3 population in RPL patients compared to healthy controls, affirming the projection accuracy of ProjectSVR (Fig. 3E, F). Taken together, this analysis showed that ProjectSVR enables the successful projection of query cells even with transcriptome bias affected by either non-biological (stress response) or biological (disease) onto the reference atlas, and facilitates the interpretation of cell population change in the disease-control design scenario.

**Figure 3.**
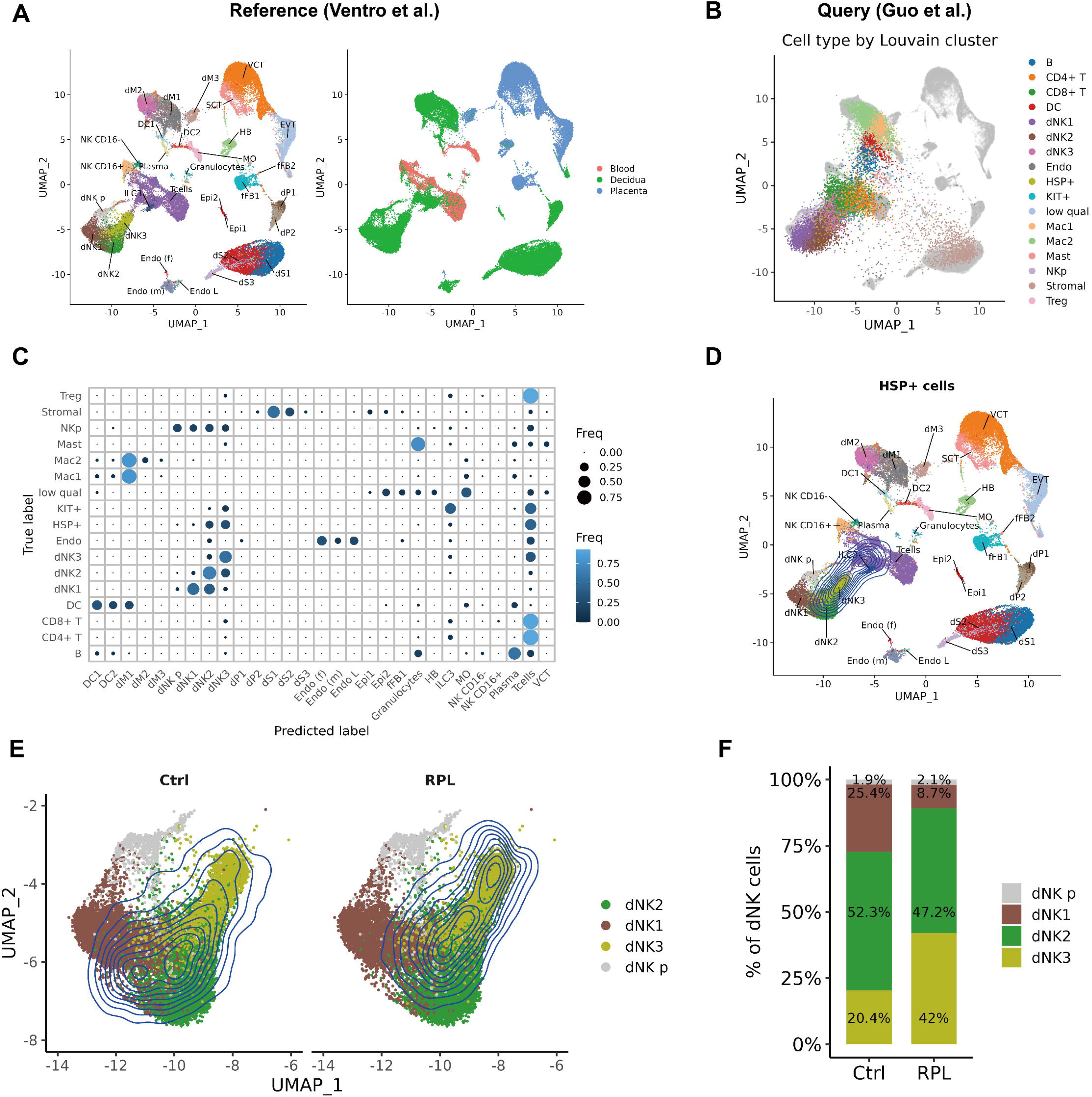
Projection analysis of decidual immune cells between recurrent pregnancy loss (RPL) patients and healthy controls. We built a cell atlas of maternal-fetal interface from Ventro et al (2018) and mapped decidual immune cells of RPL patients. (A) UMAP plots showing the cell atlas of maternal-fetal interface. Each dot represents a cell colored by cell type (left) or tissue (right). The cell labels were obtained from the original study. (B) Projection of decidual cells (query) from RPL patients and controls onto reference (A). Grey dots represent the reference, query cells are colored by manually annotated cell types using Louvain cluster on harmony embeddings (see also Figure S3C). (C) Dot plot comparing true cell labels (columns), defined by clusters (B), to predicted cell labels (rows) from reference (A) using ProjectSVR. The dot size represents the frequency of predicted labels in each true label (the sum of rows is normalized to 1). (D) 2D density plot showing the projection of cells highly expressing heat shock proteins (HSP+ cells) onto the reference. Each dot represents a reference cell, colored by reference cell type. (E) Density of the predicted decidual NK (dNK) cells from controls and RPL patients mapped to dNK reference. (F) The distribution of dNK cells from controls and RPL patients corresponding to panel (E).

### ProjectSVR enables accurate identification of the tumor-infiltrated T cell heterogeneity in cancer immunology study

The complexity and plasticity of T cells pose challenges for studying adaptive responses in the context of cancer ^13^. The recently published pan-cancer T cell atlas presents a comprehensive annotation of heterogenous T cell states ^15^. The authors employed a complex integration strategy to aggregat 390,000 T cells from 316 patients with 21 different types of cancer into two harmonious “meta cell” atlases. This T cell lanscape offers an opportunity to consistently define T cell states in studies on the tumor-infiltrated T cells. However, the integration methods used in this atlas prevent direct projection of query data using commonly employed reference mapping algorithms ^12, 13, 23^. To address this issue, we utilized ProjectSVR on the pan-cancer CD4 and CD8 T cell atlas. We trained a regression model using gene set scores calculated from the cluster-specific genes identified by the authors. Subsequently, we projected a dataset of breast cancer cases classified as either responders or non-responders to anti-PD1 treatment in order to investigate differences in the heterogeneity of tumor-infiltrated T cells between these two groups. Consistent with the original study ^24^, we observed that T helper (Tfh) cells, T helper 1 (Th1) cells (referred to “experienced T cells” in the original study) and regulatory T cells (Treg) were enriched in the pretreatment tumors of responders (Fig. 4A). The differential analysis of CD4 T cell components indicated that IFNG+ Tfh/Th1 (c17) and IL21+ Tfh cells (c16) were the top two enriched clusters ordered by p-value in anti-PD1 treatment responders (Fig. 4B and C). Furthermore, reference mapping and k-NN label transfer revealed a significant increase in the proportion of GZMK+ (c11) and terminal (c12) exhausted CD8 T cells in responders (Fig. 4D-F). These findings align with the conclusions drawn in the original study ^24^.

**Figure 4.**
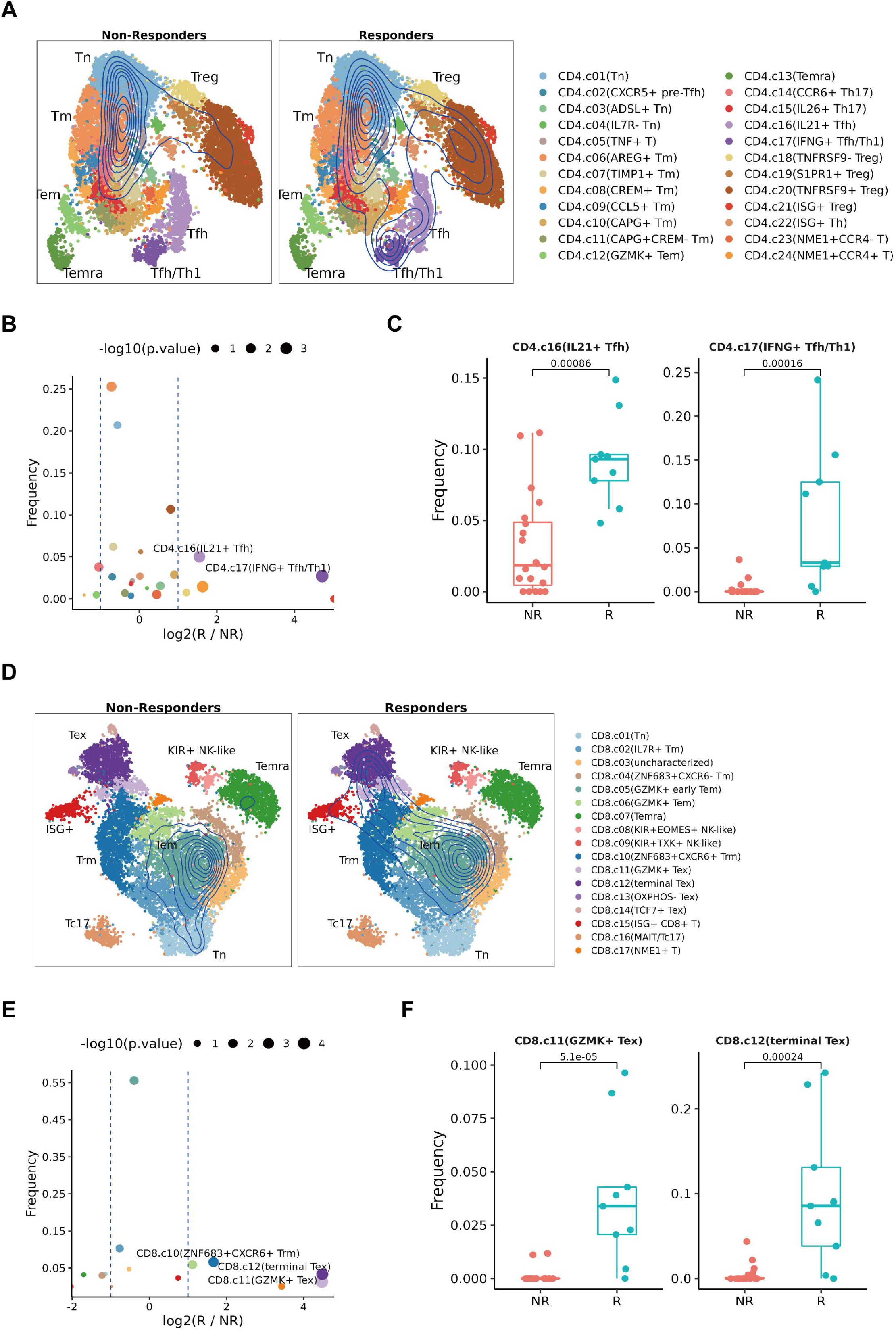
Interpreting the tumor-infiltrated T cells via ProjectSVR. We projected tumor-infiltrated CD4+ (A-C) and CD8+ T (D-F) cells from patients with breast cancer responding or non-responding to anti-PD1 treatment on the pan-cancer T cell atlas using ProjectSVR. And then transferred the reference T cell subtypes to query cells using a 10-NN classifier on reference embeddings. (A) Projection of CD4+ T cells from responders and non-responders of anti-PD1 treatment by ProjectSVR. Dots represent reference metacells colored by cell type obtained from the original study. (B) Volcano plot of differential cell subtypes of CD4+ T cells between responders (R) and non-responders (NR). The P values were calculated by Wilcoxon test. Dots colored by cell subtypes are shown in (A). (C) Boxplots comparing the frequency of two CD4+T cell subtypes between responders (R) and non-responders (NR). The P values by Wilcoxon test are shown. (D-F) The same plot as in (A-C) was applied to CD8+ T cells. (E) Dots colored by cell subtypes are shown in (D). Tfh, follicular helper T cells; Th1, T helper 1 cells. Treg, regulatory T cells. Tex, exhausted T cells; ISG, interferon-stimulated genes; Temra, terminally differentiated effector memory or effector; Tem, effector memory T cells; Trm, tissue-resident memory T cells; Tn, naïve T cells; and KIR, killer cell immunoglobulin-like receptors.

As another example, we conducted a reanalysis of the data on PD1-antibody treatment for melanoma originallyh presented in the study by Feldman et al ^25^. In agreement with the previous analysis, our findings revealed that ICB-responsive tumors exhibited a higher proportion of naïve CD8 T cells (c01) and a lower proportion of terminal exhausted CD8 T cells (c12) (Fig. S4). Additionally, we identified a rare population of Tc17 cells that was elevated in the responsive group, as reported ^15^ (Fig. S4).

Hence, these two examples provide further evidence of the efficacy of ProjectSVR in accurately projecting and characterizing T cell states in cancer immunology studies.

### Mapping the genetically perturbed germ cells to mouse testicular cell atlas

The accumulation of public scRNA-seq datasets and advances in data integration algorithm makes constructing an organ scale atlas possible ^3^. The organ scale atlas provides a baseline reference for future scRNA-seq analysis in the field of interest. In this study, we built a mouse testicular cell atlas (mTCA) to test whether ProjectSVR is suitable for reference mapping at the organ-scale atlas.

We collected mouse testicular cells from nine scRNA-seq datasets, spanning from E6.5 embryos to adults (Fig. S5A, B, and Table S2), and integrated them via the scVI algorithm implemented in scvi-tools ^18^. To annotate the cell types, we obtained publicly available cell type labels and harmonized them manually. Then we trained an SVM model using gene module scores calculated from cNMF-derived featured gene sets on the labeled cells to predict the types of unlabeled cells (Methods and supplementary materials). To refine the predicted cell type annotations, we employed a majority voting procedure on clusters identified on the latent space provided by scVI, using the Leiden algorithm ^26^ (Fig. S5C, methods). Finally, we constructed an mTCA comprising 34 cell types, encompassing 17 germ cell types across five major developmental stages (PGCs, prospermatogonia, spermatogonia, spermatocytes, and spermatids) as well as 12 somatic cell types (Fig. 5A, B). The germ cells represented a continuous trajectory on the UMAP embeddings, indicating their progression along a lengthy developmental process within a single lineage (Fig. 5A).

**Figure 5.**
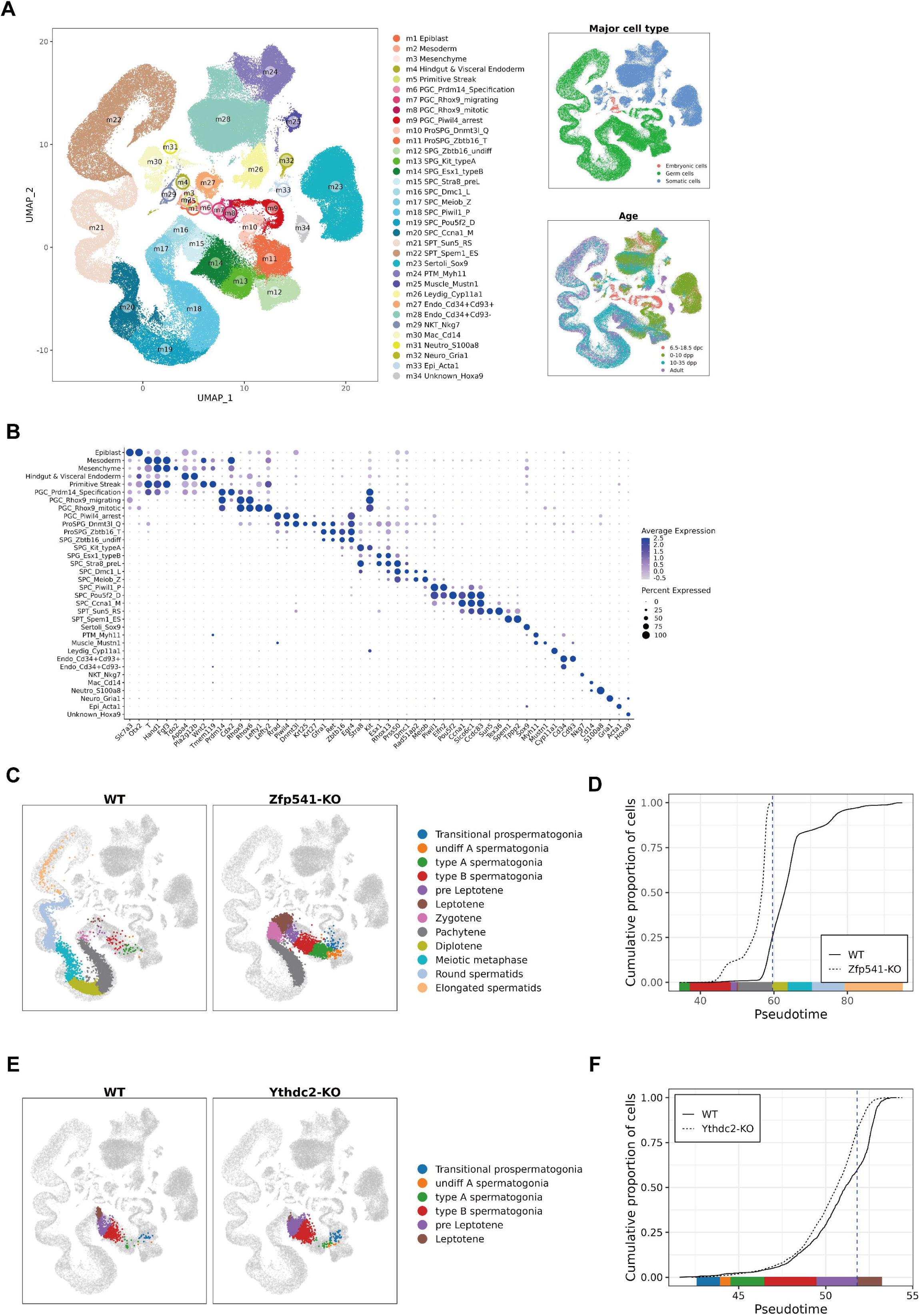
Mapping the genetically perturbed germ cells to mouse testicular cell atlas (mTCA). We built a reference of wild-type (WT) mouse testicular cells from 6.5 days post coitum (dpc) to adults and then mapped two datasets consisting of paired samples from WT and knock-out (KO) mice. (A) UMAP of WT mouse testis reference (n = 188,862 cells), colored by major cell types (upper right), age (lower right), and cell subtypes (left). Each dot represents a single cell. (B) Dot plot for expression of representative signature genes of testicular cells. The dot size represents the percentage of cells expressing the indicated genes in each stage and the dot color represents the average expression level of the genes. (C) Projection of WT and Zfp541-KO germ cells onto the testis reference. Grey dots represent reference cells, query cells colored by cell types predicted via ProjectSVR using a 10-NN classifier on reference embeddings. (D) The cumulative distribution of cells along pseudotime from each mouse strain. The pseudotime was predicted via ProjectSVR from mTCA. (E, F) The same plots as in (C, D) were applied to Ythdc2-KO germ cells.

To evaluate the capability of ProjectSVR in facilitating the analysis of new datasets in the context of mTCA, we mapped query data comprising 17,253 germ cells obtained from WT and *Zfp541*^-/-^ mouse testes onto mTCA ^27^. By employing a KNN classifier (k=10), we compared the predicted cell labels with the original annotations provided by the authors. The results yielded an overall accuracy of 0.758 (Fig. S6A, Table S3). Additionally, we transferred the latent pseudotime of the mTCA germ cells to the query cells and showed a strong correlation with the independently calculated pseudotime values by the authors (r = 0.973, Fig. S6B). This accurate prediction of cell labels and pseudotime indicated that Zfp541^-/-^ germ cells fail to progress into the diplotene stage, which is consistent to the original paper ^27^ (Fig. 5C, D).

In another example, we applied ProjectSVR to project the WT and *Ythdc2*^-/-^ germ cells collected from the 10-day postpartum (dpp) mouse testes to mTCA ^28^. Notably, we found that *Ythdc2*^-/-^ germ cells arrest at the pre-leptotene stage (Fig. 5E, F), which aligns with previously reported cytologic observations ^29, 30^. These two case studies demonstrate the efficacy of ProjectSVR in accurately and intuitively projecting a continuous developmental trajectory with transcriptional gradients onto an organ-scale reference atlas. Moreover, it facilitates the interpretation of developmental perturbations resulting from the loss of specific genes.

### Evaluating in vitro induced meiosis using ProjectSVR

Adopting the organ-scale reference atlas to evaluate the in vitro development procedure is an appealing application. In this case, we projected germ cells that were dissociated from the testis of 6-day-old mouse pups and undergoing in vitro meiosis induced by co-treatment of nutrient restriction and retinoic acid (NRRA) onto mTCA ^31^. The projected results indicated that the cultured spermatogonia differentiated along the developmental trajectory after NRRA induction (Fig. 6A). To determine the stage of the induced germ cells, we transferred the cell labels from mTCA to query cells. We observed that after two days of NRRA treatment, all of the spermatogonia differentiated. By day three, a proportion of germ cells initiated meiosis (Fig. 6B and C). Meiosis marker genes began expressing at pre-leptotene stage, while the pre-meiosis markers were repressed (Fig. 6D). These results confirmed that the NRRA treatment could induce cultured spermatogonia into meiosis in vitro without Sertoli cells, as reported in the original study^31^.

**Figure 6.**
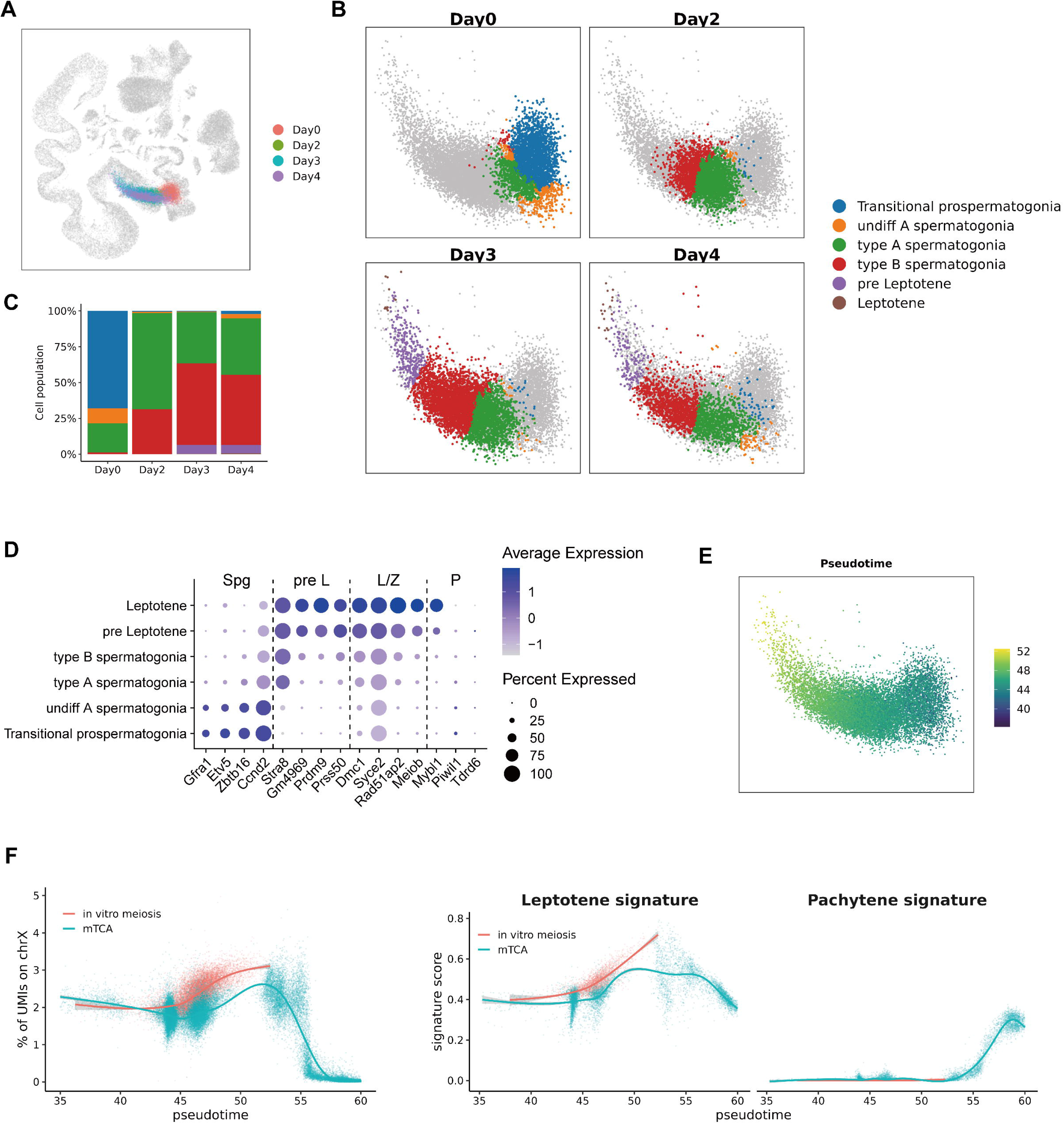
Comparing the in vitro cells to the in vivo reference. We mapped the mouse germ cells undergoing in vitro induced meiosis to mTCA built in Figure 5. (A) Projection of germ cells treated with nutrient restriction and retinoic acid (NRRA). Query cells are colored by days after NRRA treatment, and grey dots represent the reference cells in (Figure 5A). (B) Cell labels transferred from a reference in (Figure 5A) using a 10-NN classifier on reference embeddings. Colored dots, cells collected from the indicated day after NRRA treatment colored by predicted cell labels; grey dots, cells not belonging to the indicated day. (C) The distribution of germ cell stages from different days after NRRA treatment. (D)) Dot plot for expression of representative signature genes of indicated cell types. The dot size represents the percentage of cells expressing the indicated genes in each stage and the dot color represents the average expression level of the genes. Spg, spermatogonia; pre L, pre leptotene; L, leptotene; Z, zygotene; and P, pachytene. (E) Pseudotime predicted by ProjectSVR from mTCA. (F) Pseudotime analysis provides a quantitative and high-resolution evaluation of meiotic sex chromosome inactivation (MSCI) between NRRA-induced and natural meiosis. (F) Pseudotime analysis evaluating the activity of leptotene and pachytene signature between NRRA-induced and natural meiosis.

Given that pachytene cells were not detected in the projection results, and no expression of pachytene marker genes (e.g., *Piwil1* and *Tdrd6*), we hypothesized that these germ cells failed to enter pachytene (Fig. 6B and D). To verify this, we examined meiotic sex chromosome silencing (MSCI), the pachytene hall marker along the developmental trajectory. After transferring the pseudotime from mTCA to query cells, we fitted a curve for the percentage of UMIs from chromosome X along the pseudotime grouped by in vivo and in vitro meiosis (Fig. 6E). We found that unlike the meiosis in vivo, the gene located on the X chromosome in NRRA-induced meiosis germ cells were not repressed (Fig. 6F). Furthermore, we also observed that the pachytene programs were not activated, and the leptotene programs were not repressed in the most advanced germ cells undergoing induced meiosis (Fig. 6G). These results suggest that although a proportion of pachytene-like cells were identified in the spermatocyte spreading experiment, the NRRA-induced meiosis in vitro failed to progress to the pachytene stage at the transcriptome level ^31^.

## Discussion

With the rapid development of single-cell technology and decreasing sequencing costs, scRNA-seq became a powerful tool in relevant studies including cell atlas construction or profiling the landscape cellular diversity in disease or genetic perturbation. However, the inconsistency of cell annotations and clusters by different labs hinders directly comparing the novel dataset with published data, preventing us from learning general rules underlying the data from different cohorts. On the other hand, once the pieces of data scattered across studies were integrated, this give us an atlas providing a baseline and biological context and facilitating the analysis of the novel scRNA-seq dataset. As we demonstrated in the case studies, ProjectSVR provides an unbiased way to compare datasets from different technologies, disease states, genotypes, and experimental treatments in a uniform space of cell states.

The existing methods treat the reference mapping problem as a special case of data integration between a small query data and a large, well-annotated, integrated reference. Thus these methods adopt the same mathematical framework from integration to mapping. However, the mapping models are not available for many published well-constructed cell atlas, like the pan-cancer tumor-inflated T cell landscape, because of a user-defined integration strategy or lack of mapping interface for many integration algorithms. This limited us to exploring query datasets via reference mapping. On the contrary, ProjectSVR treated mapping as regression, thus we can train a regression model for mapping and directly predicting the reference embeddings for the query cells, only if the digital expression matrix and final embeddings were available. In this lack-of-mapping-interface scenario, ProjectSVR may be a good choice.

Feature Engineering is a crucial step in machine learning. This process makes the model more robust, saves training time costs, avoids the curse of dimensionality, and enhances generalization ^9^. To reduce the influence of the sparsity of single-cell data on the regression model, we treated cell type-specific genes as a meta-gene and evaluated its activity through the AUCell algorithm returning gene activity scores from 0 to 1. This value is based on the rank of gene expression levels, unaffected by normalization methods, broadening ProjectSVR’s utility in mapping datasets that lack raw counts matrix. There are many choices for identifying cell type-specific genes, we implemented an over-expressed test based on clusters (suitable for discrete cell types) and matrix factorization based on cNMF (ideal for continuous cell states). This process introduces a hyperparameter: gene set size. The choice of gene set size is robust to mapping results (data not shown). Usually, a default setting (top 25 genes) used in most case studies is good enough.

In all cases, the UMAP was adopted for the final representation of reference atlases. In theory, any biological meaningful representation of dimensionality reduction is accepted for ProjectSVR. We choose UMAP in our case studies because UMAP preserves more global data structure than tSNE while keeping as much as a local data structure, which makes UMAP generate a representation that preserves the continuity of the cell subsets ^32^. Pseodotime is another intuitive representation of a continuous biological process mapping all cells on one dimension. In the case study of mTCA, we showed that ProjectSVR was able to place the query germ cells onto the correct position along the pseudotime (Figure S6B). In some cases (such as the *Ythdc2*-KO study), test cell distribution on pseudotime may be more sensitive to developmental arrest.

We expect to generate more well-annotated reference models from diverse biological systems instead of a unified single atlas for all cell types such as MCA (mouse cell atlas)^33^. To promote the reproducibility of scRNA-seq data analysis, we shared five pre-built models as the initial release of ProjectSVR. We aim to build a comprehensive database to store more reference models and provide easy-use API on a web server in further work.

## Methods

### ProjectSVR implementation details

ProjectSVR is a modularized R package for reference mapping based on supported vector regression, including feature extraction, feature scoring, model training, reference mapping, and label transfer. For any pre-trained model in hand, users can start from the reference mapping step by providing a query matrix in raw counts or normalized form. We shared all pre-trained models at https://zenodo.org/record/8191559.

### Feature extraction

ProjectSVR adopts two methods for feature selection. If clusters or cell types are provided, the marker-based method is preferred. For the atlas describing a continuum of cell states, we provide R wrappers (FindOptimalK and RunCNMF) for the python version cNMF through the R package reticulate. The high-weighted genes in each program are selected for feature scoring. To reduce the running time for the large dataset, we also provide a set of wrappers (EstimateKnnDensity, subset, and MergeCells) to merge adjacent cells into meta cells to produce a meta cell matrix for downstream cNMF.

### Feature scoring

Feature scoring quantifies the activity of the genes in a given gene set reflecting the cell types, states, or biological processes. Here, we used UCell package, a multithreading version of the AUCell algorithm to speed up the feature scoring step. UCell scores depend only on the relative expression of genes in each cell and are not dependent on dataset composition. Therefore, we can calculate UCell score from any form of gene expression matrix (raw counts, normalized or scaled matrix), which expands the usage of our model.

### Model training & Reference mapping

We test all regression algorithms implemented in the *mlr3learners* package and find SVR performs best, perhaps due to its good performance on nonlinear regression (data not shown). Considering SVR is suitable for small samples and not scalable to large samples due to the O(n^2^) time complexity, we introduced an ensemble SVR model for predicting the embedding coordinates. Before model training, L2 normalization is performed on the feature score matrix across cells. Then we randomly subsampled M cells from the input matrix with N times and train N SVR models. The final predicted coordinates of each dimension are the median value of predictions from the N models. A balanced mode was also implemented for the subsampling procedure to avoid the model bias to any cell types. The model training and reference mapping of each model are processed in parallel using the *parallel* package to reduce time consumption. We provided two functions FitEnsembleSVM and ProjectNewdata for model training and reference mapping.

### Projection metric

We defined the mean KNN distance to measure the mapping quality. The main idea is that a good mapping procedure should keep the local topological relationship after projection. We construct a KNN network in the feature space using the UCell score and calculated the average distance of the K neighbors in reference space for each cell. The mean distance of K randomly selected cells was calculated and repeated serval times to construct the null distribution. Then empirical p values were estimated using this null distribution. The p values were corrected using the Benjamini-Hochberg Procedure. A wrapper (AddProjQual) is provided for this calculation.

### Label transfer

Once query cells are projected in the same embeddings as the reference, reference labels can transfer to query cells using any classification model. We use a simple k-NN classifier and provide a convenient wrapper (*KnnLabelTransfer*), which returns the predicted labels and the proportion of reference neighbors supporting the majority vote.

### Model sharing

The reference atlases used in this manuscript are available at https://zenodo.org/record/8191576. All models trained and used in this paper are available at https://zenodo.org/record/8191559. We have provided step-by-step guides for training new models and case studies of ProjectSVR, which can be found in ProjectSVR GitHub repository (https://jarninggau.github.io/ProjectSVR/).

## Building mouse testicular cell atlas

### Data collection and preprocessing

As different pipelines generated the released mouse testicular scRNA-seq gene expression matrixes with different versions of genome assemblies and gene annotations, the FASTQ or BAM files from SRA or ArrayExpress database were downloaded and preprocessed to harmonize the count matrixes of different datasets. All reads were aligned and quantified using the STARsolo pipeline implemented in STAR (v2.7.9a) against the mm10 mouse reference genome download from 10x Genomics official website. The raw counts were used for downstream analysis. Quality control was applied to cells for each dataset. For the dataset generated by the 10x platform, we dropped the cells whose detected genes were less than 500 or the proportion of mitochondrial gene counts more than 15%. For the STRT-seq datasets, cells with detected genes less than 1000 or a proportion of mitochondrial gene counts of more than 10% were filtered out. To identify potential doublets in the 10x datasets, we run DoubletFinder (v2.0.3) by sample. The expected doublet rate was set to be 0.075, and the cells labeled ‘Singlet’ were kept. For the STRT-seq datasets, we downloaded the high-quality barcodes from the original studies, and only the cells with a matched high-quality barcode were kept. The contamination rates were calculated using decontX function implemented in the R package celda for each sample.

### Clustering per dataset

We perform clustering on each dataset to identify the contamination clusters ignored by DobuletFinder in the 10x datasets. Cells from different samples were normalized to log(CPM/100+1) and then integrated into MNN embeddings using the RunFastMNN function implemented in SeuratWrappers (v.0.3.0) with default settings according to the Seurat tutorial. The first 30 MNN dimensions were used for clustering and UMAP embedding. Then we identified contaminated clusters by examining the clustering results on the UMAP plot. The potential contamination clusters expressed a mixture of markers of multiple spermatogenesis stages and were removed from the following analysis.

### Data integration across studies

All filtered matrixes from different studies were merged into one count matrix. In total 210,415 cells were kept at this step. We used sceasy to transform formats between Seurat and h5ad files. To remove the batch effect between studies while keeping the biological variance, the scVI algorithm implemented in scvi-tools (0.17.4) was chosen for data integration as in previous reports ^34, 35^. The top 2,000 high-variable genes across datasets were identified using the highly_variable_genes function implanted in scanpy for the following analysis. Then we took the counts matrix of the 2,000 genes as input of scVI using 30 dimensions of latent spaces and 2-layer neuron networks to fit our data as described in the documentation (https://scvi-tools.org/). UMAP visualization and Leiden clustering were performed on 30 scVI latent embeddings with default parameters.

### Cell type annotation

#### Collection of public cell labels

The published cell labels were collected from original studies if available. Then we manually harmonized the labels. Besides, we manually annotated a part of the datasets according to the markers in the original studies. In total, we annotated 104,669 cells.

#### Label transfer via machine learning

To transfer these labels to the whole atlas, we trained machine learning models to predict the rest unlabeled cells. To construct features in an unsupervised way, we identified gene expression programs via consensus non-negative matrix factorization (cNMF). To reduce the running time of cNMF, we pooled the cells into metacells for each dataset using R package metacell and run cNMF on raw counts matrix of metacells using the 2000 high variable genes. We chose 100 iterations of the NMF procedure and selected a solution with 70 programs based on the elbow criterion of reconstruction error of the input data matrix and clustering stability. We extracted the top 100 weighted genes in each program and calculated the gene module score for each cell using the R package AUCell. To predict the cell types on the unlabeled data, we trained random forest, xgboost, and SVM models on the labeled data and evaluate the performance via accuracy and area under the receiver operating characteristic curve (AUROC) through 10-fold cross-validation. The SVM performed best on this dataset and was selected as the final prediction model.

#### Manual correction of predicted labels

We found that there were some clusters with very low predictive probabilities, which may be novel cell types excluded in the training dataset. We labeled these clusters as unknown and annotated them according to their representative markers. Finally, we identified muscle cells, neurocytes, epithelial cells, NK/T cells, neutrophils, and an unknown cell cluster expressing Hoxa9.

### Finetune the final reference

To improve the data quality, we further removed cells with high contamination rates (> 0.4). The dataset GSE113293 was removed due to its lower sequence depth compared to other datasets. Then we repeated the data integration step using scVI tools and finetune the data representation via the scANVI algorithm, which takes into account the cell identity. The 30 scANVI latent spaces were used to generate the UMAP embeddings in Figure 5A.

The prediction was performed on each cell independently. To combine the cell type predictions with the cell-cell transcriptome relationships, we adopted an iterative cluster procedure. For each epoch of clustering, we adopted the Leiden algorithm with a lower granularity at resolution = 0.1, then the cluster entropy was calculated for each cluster as 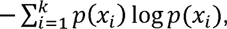, where *x_1_*,…, *x_k_* are the set of manually corrected cell labels, and *p*(*x*_i_) is the fraction of cells labeled as *<u>x</u>_i_* in the cluster. The Leiden clustering was performed on the clusters with high entropy until the given cluster was purified. The final clusters were assigned to the identity supported by the dominant manually corrected cell types.

To accommodate labels at different granularity, we made three-level hierarchical cell type annotations as described. The level1 includes the coarsest labels (germ cells, somatic cells, etc.), and level3 contains the finest labels (e.g. leptotene spermatocytes). To generate the cell markers of each cell type, we downsampled each cell type to 200 cells and calculated representative genes using the FindAllMarkers function implemented in Seurat.

### Calculation of germ cell time

For continuous biological processes such as spermatogenesis, a continuous representation is a better choice to describe a developmental spectrum ^36^. We embedded the germ cells of postnatal mice into a two-dimensional diffusion map using 30 scANVI latent spaces and calculated the pseudotime through R package destiny. The pseudotime dependent on the transcriptome difference and the cell number does not reflect the real development time. Thus, to make the pseudotime distributed more equally, we weighted the spermatogonia, spermatocytes, and spermatids equally and rescale the pseudotime on the original outputs.

## Case study

### Analysis of 10x PBMCs

#### Constructing the PBMCs query assayed by different 10x protocols

The PBMCs query datasets were each sequenced with different 10x protocols: 3’v1 (n = 4809 cells), 3’v2 (n = 8380 cells), and 5’ (n = 7697 cells). The raw count matrix and cell labels were obtained from the Harmony publication.

#### Constructing the PBMCs reference

We downloaded the pre-built blood atlas from the DISCO database. The blood atlas consists of 167,594 well-annotated PBMCs from 100 samples with different disease states. The original dataset was not included UMAP embeddings, so we integrated these cells into harmonized UMAP embeddings. We treated the cells from different studies as different batches, then we adopted the scVI tools pipeline with the same parameters as described in the *data integration* part of building mTCA. The cells labeled as ‘TRDV2 gdT’ were removed due to their scattered arrangement on the UMAP plot.

#### Mapping query cells onto PBMCs reference

The raw count matrix of PBMCs reference was first normalized by log(CP10K+1). Then the log normalized matrix was downsampled to 200 cells per cell subtype. The mitochondria and ribosome genes were excluded from marker selection. The positive markers for each cell subtype in PBMCs reference were calculated and arranged by adjusted p values using the Wilcoxon test on the downsampled log normalized matrix and the top 25 markers of each cell subtype were selected as features. Then the signature matrix was obtained from the raw count matrix using the R UCell package and L2 normalized. Then we trained the ensemble SVR models to predict the UMAP embeddings from the signature matrix using the FitEnsembleSVM function in the ProjectSVR package with 40 submodels and 5000 cells for each submodel. To map the query onto scVI integrated reference, we transformed the raw count matrix of query cells into a signature matrix as mentioned above, and took it as input for the ProjectNewdata function. The median values of the predicted reference embeddings of 40 submodels were the final outputs.

#### Benchmark to the de novo integrated embeddings

The raw count matrixes of three query datasets were merged and normalized to log(CP10K+1). Then top 2000 high variable genes were calculated using the variance stabilizing transformation (VST) procedure on the merge normalized matrix. We ran harmony on the top 30 principle components (PCs) and corrected batch effects from protocols with default settings. UMAP was performed on 30 original PCs (no correction) or corrected PCs (harmony correction). To evaluate the sample mixing between de novo integration and mapping, we calculated the Local Inverse Simpson Index (LISI) using the compute_lisi function from the R package LISI with default parameters.

#### Evaluate the accuracy of cell type assignment

We transfer the labels of cell subtypes from PBMCs reference to query cells using the KNNLabelTransfer function defined in the ProjectSVR package. Briefly, we calculated K nearest reference cells for each query cell (K = 10) using the nn2 function in the “RANN” R package and assign the cell labels supported by major neighbors. We calculated the specificity and sensitivity of each cell subtype. Specificity = TP/(TP+FP), sensitivity = TP/(TP+FN).

### Projecting decidual immune cells from RPL patients to maternal-fetal interface cell atlas

#### Building the healthy maternal-fetal interface cell atlas

The maternal-fetal interface reference datasets along with author-defined cell clusters and labels (Ventro et al.; 10x, n = 64,734 cells; smart-seq2, n = 5,591 cells) were downloaded from ArrayExpress database under E-MTAB-6678 and E-MTAB-6701. The raw count matrixes from different technologies were normalized to log(CP10K+1). Then top 2000 high variable genes were calculated using the VST procedure on cells from each technology and pooled together. The 10x and smart-seq2 datasets were integrated using the mutual nearest neighbors (MNN) algorithm implemented in the Seurat package using the RunFastMNN function with default settings. The top 30 MNN embeddings were used for UMAP visualization with default parameters.

#### Construct RPL query dataset

The RPL decidual immune dataset (Guo et al.) includes two RPL patients (n = 8908 cells) and three controls (n = 11,613 cells) assayed with 10x. The author did not release processed matrix and cell labels, so we downloaded the raw fastq files from the GSA database under CRA002181. STARsolo pipeline implemented in STAR (v2.7.9a) was adopted to align and quantify reads against the GRCh38 human reference genome download from 10x Genomics official website. The raw count matrixes from different samples were normalized and integrated using the wrapper of the harmony algorithm implemented in the Seurat package. The cells were clustered and annotated according to the expression levels of cell type markers used in the original study. Clusters without any known markers were removed (n = 72 cells). We identified a cluster expressing high heat shock proteins (HSP) and named it HSP+ cells, which were not shown in the original study.

#### Mapping query cells onto MNN integrated reference

The signature matrix was calculated via the UCell package using the top 25 most significant positive genes identified from the reference for each cluster. We projected the query cells to the reference using the FitEnsembleSVM and ProjectNewData function implemented in the ProjectSVR package with default settings described in the 10x PBMC task. The labels of cell types from the maternal-fetal interface reference defined by the original study were transferred to query cells using the KNNLabelTransfer function in the ProjectSVR package with defined settings (k = 10).

### Immune checkpoint blockade examples

#### Building the tumor-infiltrated T cell reference model

The normalized matrix, cell labels, UMAP embeddings, and cluster-specific markers of the pan-cancer T cell atlas were downloaded from https://zenodo.org/record/5461803 (CD4 T cells, n = 10,621 metacells; CD8 T cells, n = 11,972 metacells). The top 20 most significant cluster-specific markers were selected to build the gene sets for calculating the signature matrix via the UCell package. The reference model was trained using the FitEnsemblSVR function in the ProjectSVR package with defined settings (batch.size = 5000, n.models = 50).

#### Exploration of the T cell heterogeneity in ICB-responsive breast cancer and melanoma

The raw count matrix and author-defined cell labels of the breast cancer dataset were downloaded from https://lambrechtslab.sites.vib.be/en/single-cell. The T cells were extracted according to the author-defined cell labels. The normalized matrix and samples’ metadata of the melanoma T cell dataset were downloaded from the GEO database under GSE120575. We project the query CD4 and CD8 T cells to the corresponding reference using the ProjectNewData function with default settings. The cell labels from the reference T cell atlas were transferred to query cells using the KNNLabelTransfer function with defined parameters (k = 10). Wilcoxon test was performed to estimate the p values of the differential distributed T cell states between ICB responsive and non-responsive groups.

### Mouse testis examples

#### Building the mTCA reference model

To identify gene expression programs of cell-type identity, we performed cNMF on mTCA. We first grouped the adjacent cells on UMAP spaces into meta cells using EstimateKnnDensity and MergeCells functions in the ProjectSVR package to reduce the running time of cNMF. We run 50 iterations of the NMF procedure on the normalized meta-cell matrix and chose 70 NMF components based on the elbow criterion of reconstruction error of the input matrix and clustering stability through FindOptimalK and RunCNMF function in ProjectSVR package. Then top 100 weighted genes in each component were selected as cell identity markers for the computation of the gene module score using AUCell. We kept 49 cNMF components related to specific cell types, including 18 components related to germ cells for model training. We trained two integrated SVR models on UMAP embeddings of all cells (UMAP model, all 49 components) and pseudotime of germ cells (germ cell time model, 18 germ cell-related components) respectively using the related gene module score. For the *Ythdc2*-KO example, as some gene names remained discordant, we use the intersected genes between the query dataset and mTCA reference to calculate the gene set score and train models.

#### Analysis of scRNA-seq data of Zfp541-KO and Ythdc2-KO germ cells

The raw count matrix and cell metadata of *Zfp541*-KO and *Ythdc2*-KO datasets were downloaded from the GEO database under GSE172157 and GSE196427. We mapped the query cells to mTCA reference using ProjectNewdata with default parameters. We used 10-NN to transfer reference labels to query cells using the KnnLabelTransfer function. The mapping quality was estimated using the AddProjQual function in the ProjectSVR package with defined settings (k = 10, repeats = 10000). Only projected cells with adjusted p value < 0.05 were kept for further analysis. To distinguish the stages that spermatocyte arrest, we predicted the germ cell time for the query germ cells.

#### Mapping in vitro meiosis against mTCA

The raw count matrix in h5 format was downloaded from the GEO database under GSE153274. The projection, mapping quality, and label transfer were performed described in the upper section. The leptotene and pachytene signature scores were calculated through the AUCell package using the top 100 most significant leptotene or pachytene marker genes.

## Supplementary Figure legends

**Figure S1.**
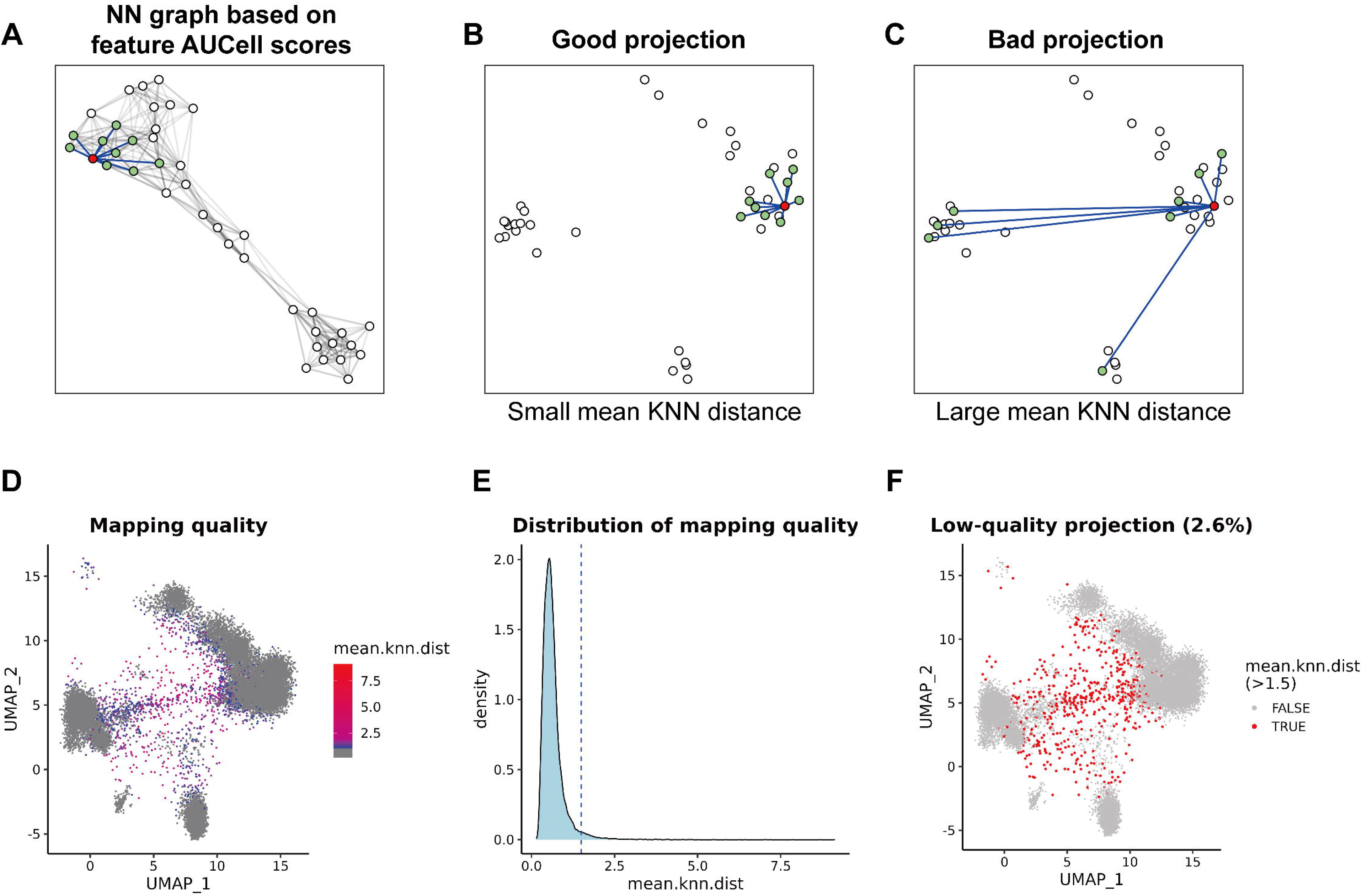
The mean K nearest neighbors (KNN) distance metric. To evaluate the mapping quality of query cells onto a reference, we defined a metric called mean KNN distance. (A) To calculate the mean KNN distance on reference embeddings, we first construct a KNN graph in feature space (UCell scores). (B, C) Then we calculated the mean distance of the k nearest neighbors of a given cell on reference embeddings. (B) If the given cell kept the local relationship of its k nearest neighbors after projection, it will have a small mean KNN distance and indicate a good projection. (C) Otherwise, a large mean KNN distance means a bad projection. (D-E) The mapping quality of the projected PBMC dataset (see also Figure 2). (D) Projected embeddings of three PBMC datasets. Dots are colored by mean KNN distance (mean.knn.dist). (E) Distribution of mean.knn.dist corresponding to panel (D). The dashed line indicating the cutoff value (1.5) of mean.knn.dist represents a bad projection. (F) Low-quality projected cells (mean.knn.dist > 1.5) on projected embeddings.

**Figure S2.**
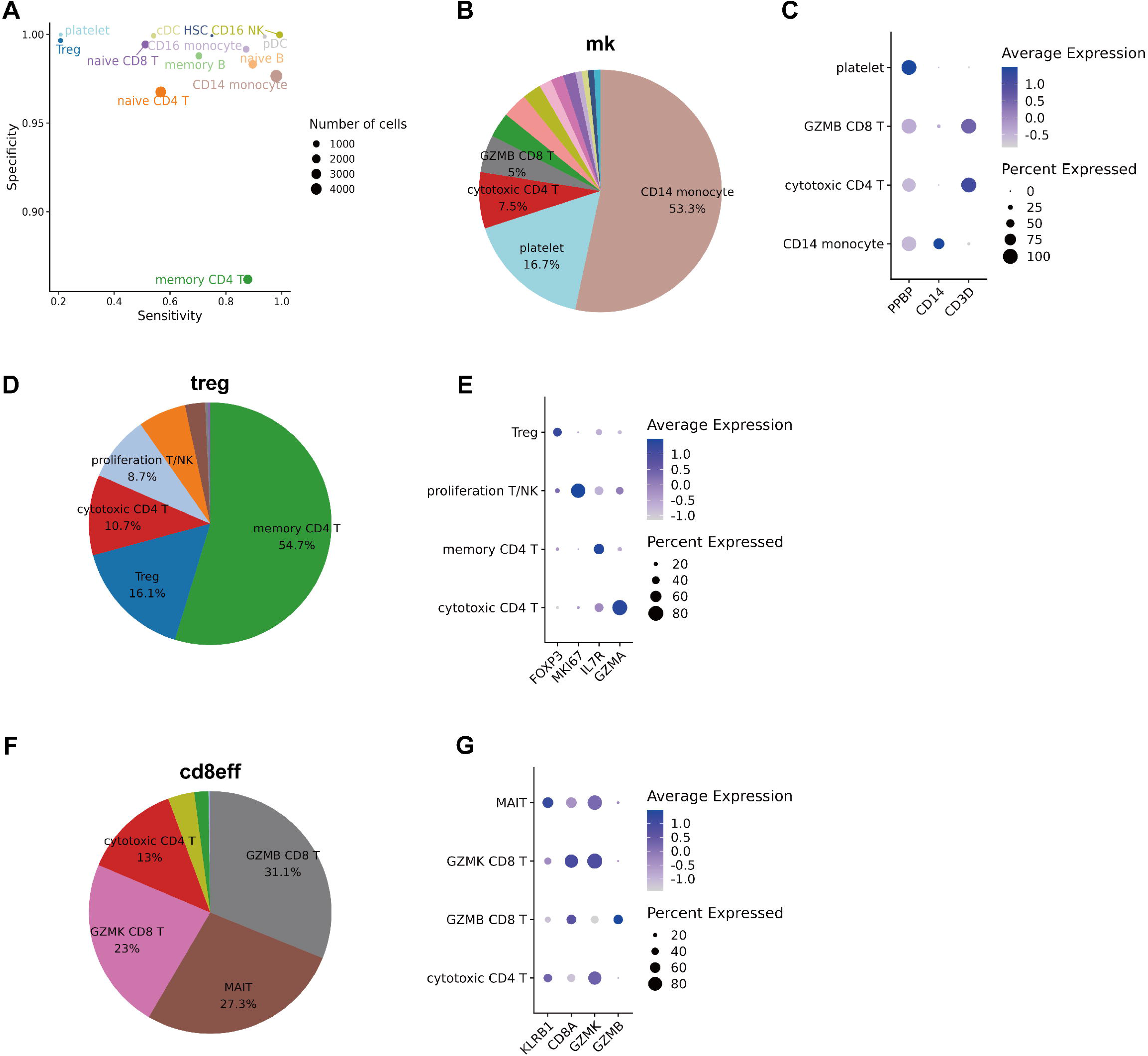
The accuracy of the predicted labels of query PBMC datasets by ProjectSVR. (A) Performance of the ProjectSVR measured by the specificity and sensitivity for query PBMC dataset. (B) Pie plot showing the distribution predicted labels in original annotated megakaryocytes(mk). (C) Dot plot for expression of representative signature genes of predicted labels from original megakaryocytes in (B). The dot size represents the percentage of cells expressing the indicated genes in each stage and the dot color represents the average expression level of the genes. (D, E) The same plots as in (B, C) applied to regulatory T cells (Treg). (F, G) The same plots as in (B, C) applied to effector CD8+ T cells (cd8eff).

**Figure S3.**
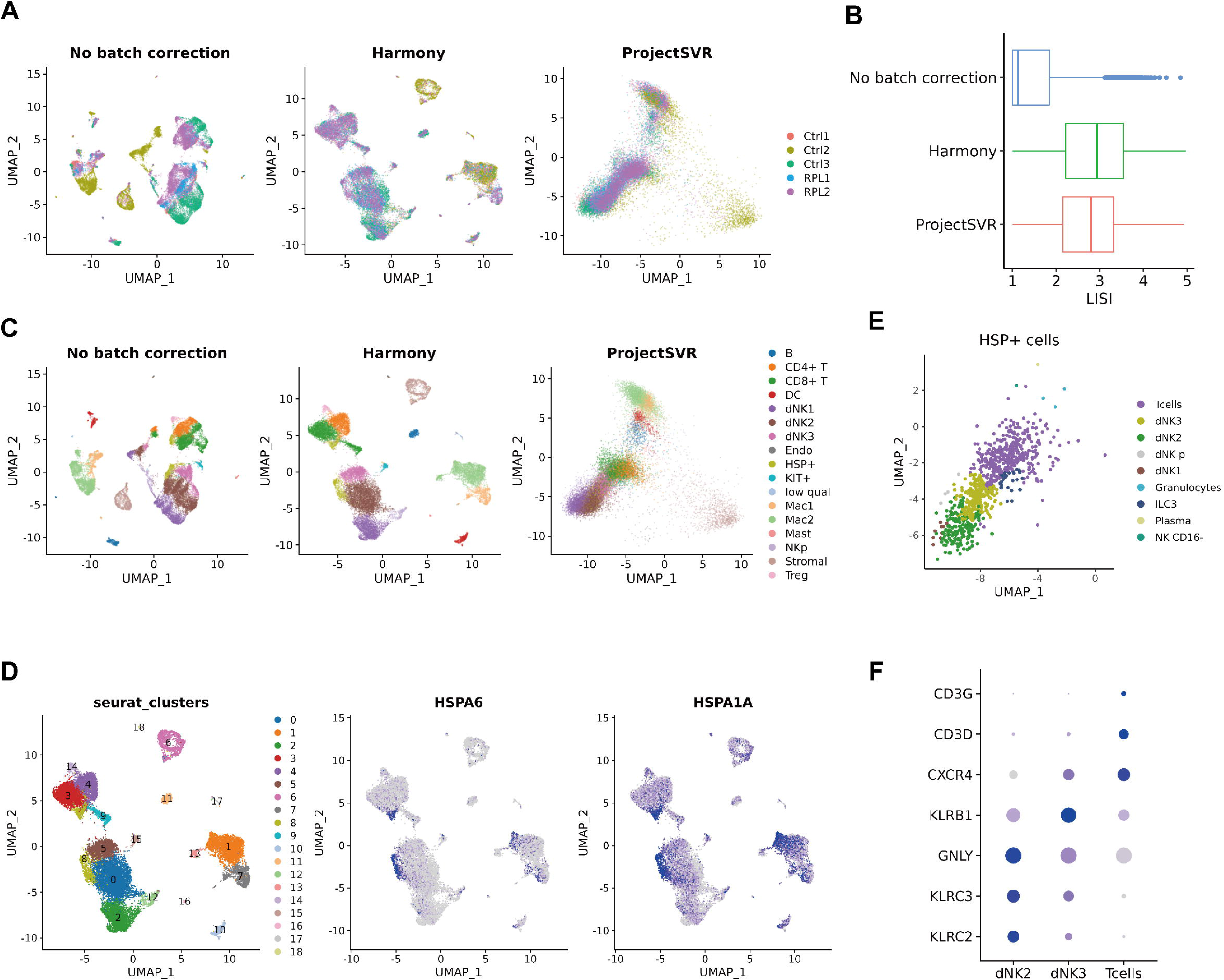
Analysis of decidual immune cells from RPL patients and controls. (A) Qualitative evaluation of batch effect correction by Harmony and ProjectSVR for decidual immune cell dataset. Dots colored by sample ID. (B) Quantitative evaluation of batch effect correction by Harmony and ProjectSVR using LISI (Local Inverse Simpson’s Index). (C) The sample plots as in (A), dots colored by cell types manually annotated using Louvain clustering on harmony embeddings. (D) Louvain clusters on harmony embeddings. Cluster 8 highly expressing heat shock proteins (e.g. HSPA6 and HSPA1A) was labeled as HSP+ cells. (F) Gene signatures of predicted cell types by ProjectSVR in HSP+ cells. The dot size represents the percentage of cells expressing the indicated genes in each stage and the dot color represents the average expression level of the genes.

**Figure S4.**
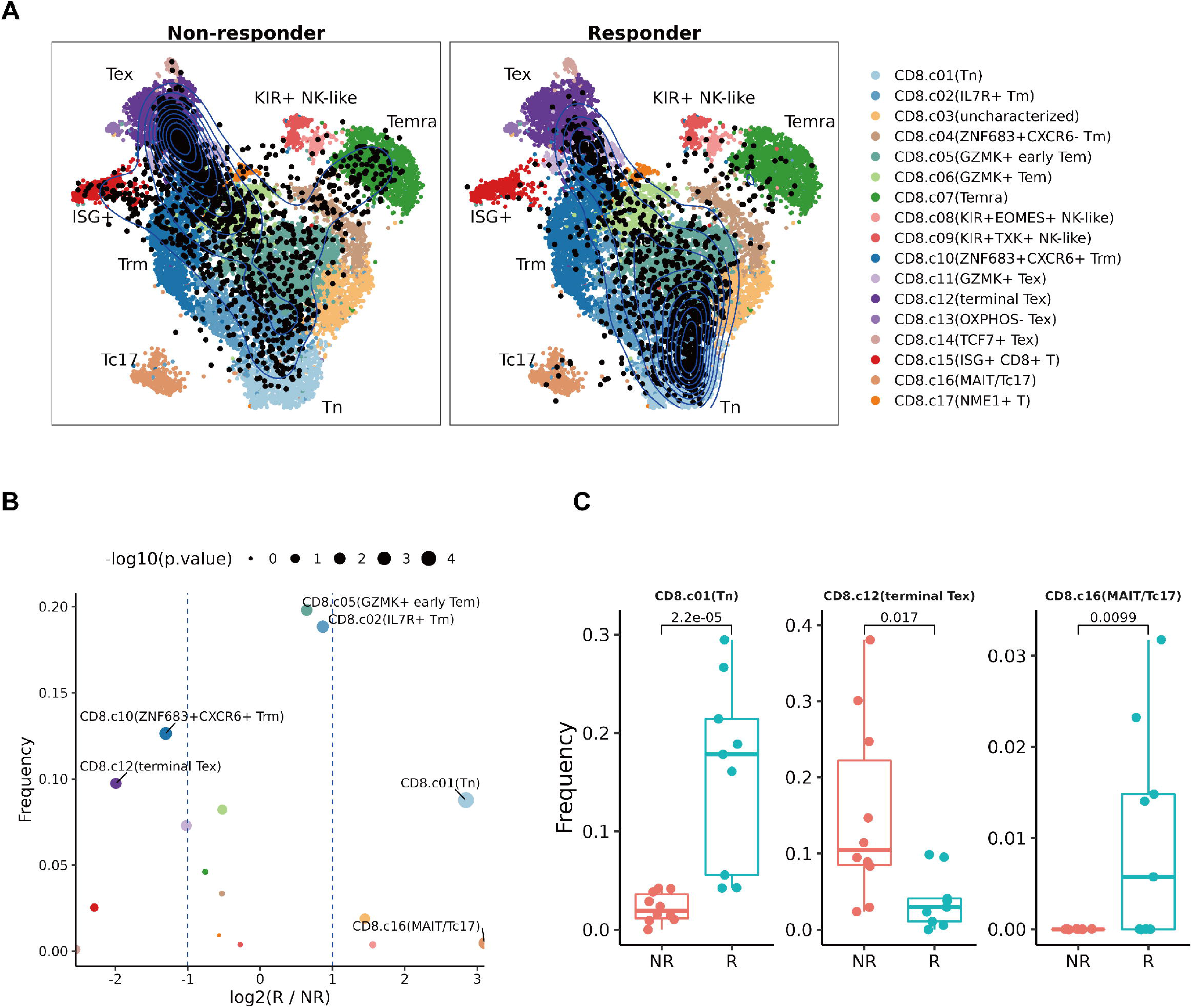
Interpreting the CD8+ T cells heterogeneity in checkpoint blockade therapy of melanoma. We re-analysis of tumor-infiltrated CD8+ T cells from melanoma patients responding and non-responding to PD1-antibody treatment (data from Feldman et al.) using ProjectSVR. We first project query cells onto pan-cancer reference, and then transferred the reference labels to query cells using a 10-NN classifier on reference embeddings. (A) Projection of tumor-infiltrated CD8+ T cells from non-responders and responders. Black dots represent query cells and colorful dots represent CD8+ T metacells form pan-cancer reference defined by Zheng et al. (B) Volcano plot of differential cell subtypes of CD8+ T cells between responders (R) and non-responders (NR). The P values were calculated by Wilcoxon test. Dots colored by cell subtypes are shown in (A). (C) Boxplots comparing the frequency of two CD8+T cell subtypes between responders (R) and non-responders (NR). The P values by Wilcoxon test are shown.

**Figure S5.**
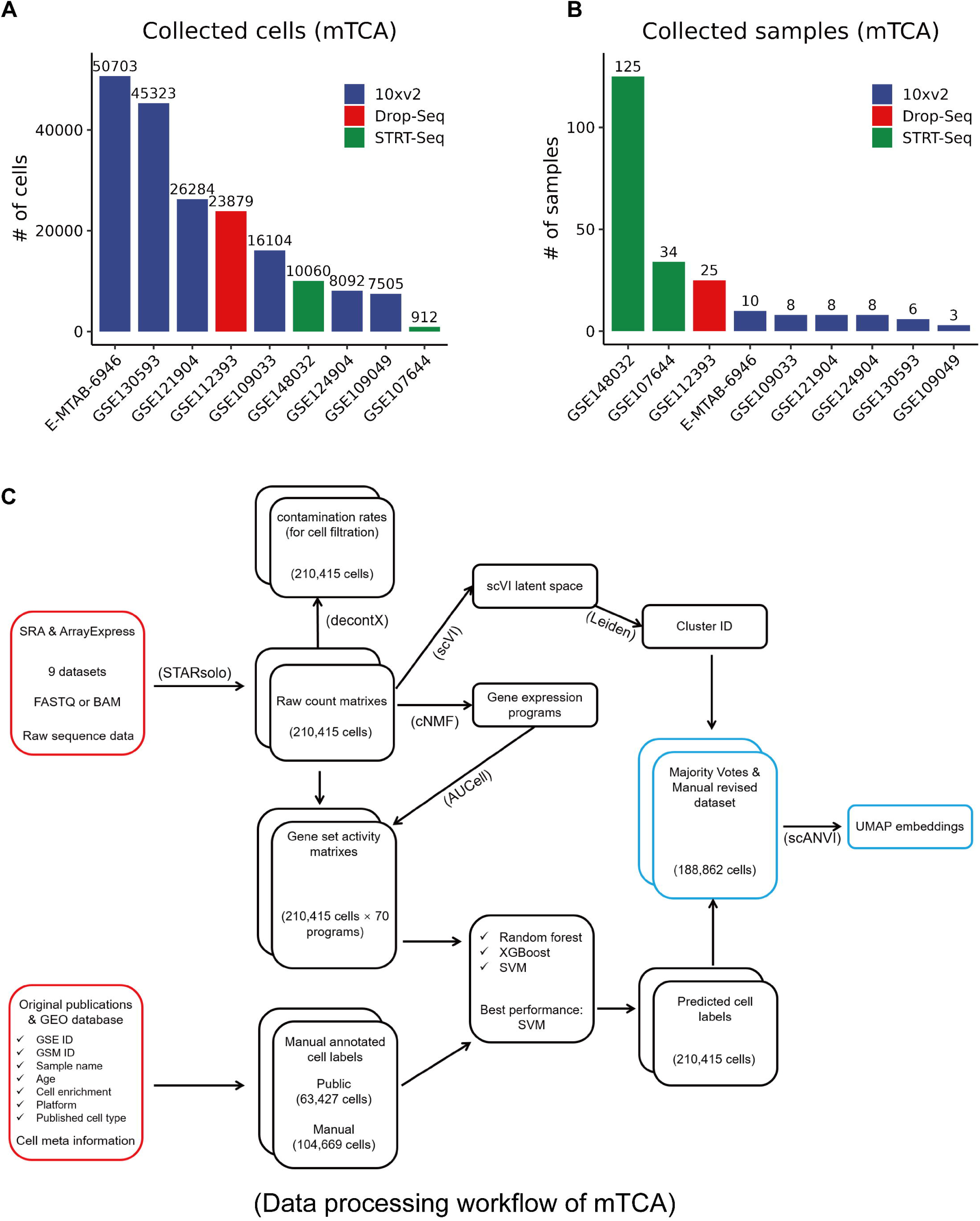
Building the mouse testicular cell atlas (mTCA). (A, B) Collection of scRNA-seq data for mouse testis. The bar plots showing the number of cells (A) and samples (B) from GEO database and ArrayExpress. (C) Schematic representation of the preprocessing and annotation steps for building mTCA.

**Figure S6.**
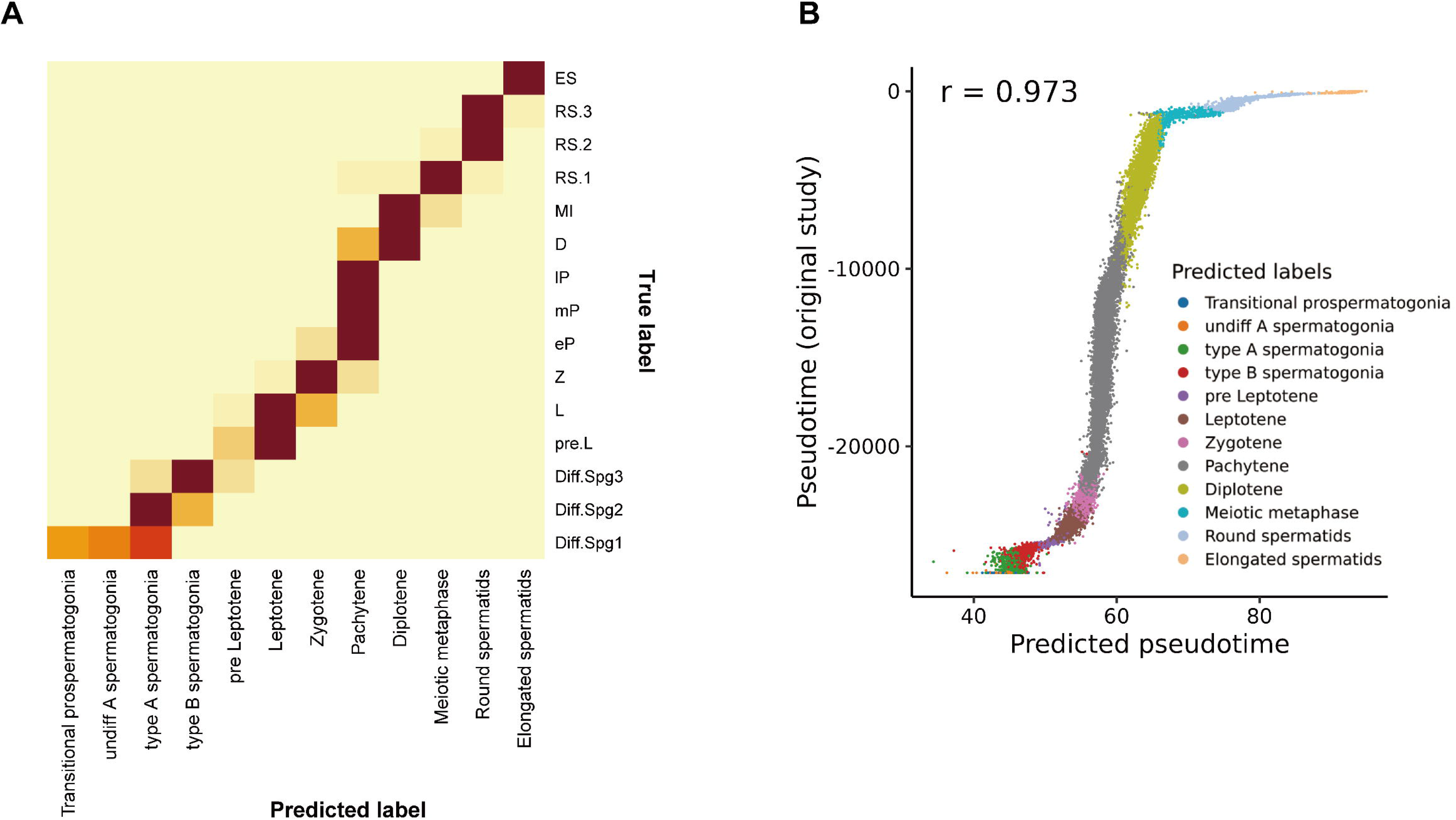
The performance of ProjectSVR in case study of Zfp541-KO germ cells. (A) Confusion matrix comparing the predicted labels and True labels. True labels were obtained from the original study by Xu et al. (B) Correlation of the pseudotime from the original study and predicted by ProjectSVR. The Spearman’s rho was shown.

## References

1. Method of the year 2013. Nat Methods 11, 1, doi:10.1038/nmeth.2801 (2014).

2 Luecken, M. D. et al. Benchmarking atlas-level data integration in single-cell genomics. Nat Methods 19, 41–50, doi:10.1038/s41592-021-01336-8 (2022).

3. Sikkema, L., et al. An integrated cell atlas of the human lung in health and disease. bioRxiv, 2022.2003.2010.483747, doi:10.1101/2022.03.10.483747 (2022).

4 Vento-Tormo, R. et al. Single-cell reconstruction of the early maternal-fetal interface in humans. Nature 563, 347–353, doi:10.1038/s41586-018-0698-6 (2018).

5 Han, X. et al. Construction of a human cell landscape at single-cell level. Nature 581, 303–309, doi:10.1038/s41586-020-2157-4 (2020).

6 Lahnemann, D. et al. Eleven grand challenges in single-cell data science. Genome Biol 21, 31, doi:10.1186/s13059-020-1926-6 (2020).

7 Abdelaal, T. et al. A comparison of automatic cell identification methods for single-cell RNA sequencing data. Genome Biol 20, 194, doi:10.1186/s13059-019-1795-z (2019).

8 Huang, Q., Liu, Y., Du, Y. & Garmire, L. X. Evaluation of Cell Type Annotation R Packages on Single-cell RNA-seq Data. Genomics Proteomics Bioinformatics 19, 267–281, doi:10.1016/j.gpb.2020.07.004 (2021).

9 Pasquini, G., Rojo Arias, J. E., Schafer, P. & Busskamp, V. Automated methods for cell type annotation on scRNA-seq data. Comput Struct Biotechnol J 19, 961–969, doi:10.1016/j.csbj.2021.01.015 (2021).

10 Galdos, F. X. et al. devCellPy is a machine learning-enabled pipeline for automated annotation of complex multilayered single-cell transcriptomic data. Nat Commun 13, 5271, doi:10.1038/s41467-022-33045-x (2022).

11 Hao, Y. et al. Integrated analysis of multimodal single-cell data. Cell 184, 3573–3587 e3529, doi:10.1016/j.cell.2021.04.048 (2021).

12 Kang, J. B. et al. Efficient and precise single-cell reference atlas mapping with Symphony. Nat Commun 12, 5890, doi:10.1038/s41467-021-25957-x (2021).

13 Andreatta, M. et al. Interpretation of T cell states from single-cell transcriptomics data using reference atlases. Nat Commun 12, 2965, doi:10.1038/s41467-021-23324-4 (2021).

14 Tran, H. T. N. et al. A benchmark of batch-effect correction methods for single-cell RNA sequencing data. Genome Biol 21, 12, doi:10.1186/s13059-019-1850-9 (2020).

15 Zheng, L. et al. Pan-cancer single-cell landscape of tumor-infiltrating T cells. Science 374, abe6474, doi:10.1126/science.abe6474 (2021).

16 Kotliar, D. et al. Identifying gene expression programs of cell-type identity and cellular activity with single-cell RNA-Seq. Elife 8, doi:10.7554/eLife.43803 (2019).

17 Andreatta, M. & Carmona, S. J. UCell: Robust and scalable single-cell gene signature scoring. Comput Struct Biotechnol J 19, 3796–3798, doi:10.1016/j.csbj.2021.06.043 (2021).

18 Lopez, R., Regier, J., Cole, M. B., Jordan, M. I. & Yosef, N. Deep generative modeling for single-cell transcriptomics. Nat Methods 15, 1053–1058, doi:10.1038/s41592-018-0229-2 (2018).

19 Li, M. et al. DISCO: a database of Deeply Integrated human Single-Cell Omics data. Nucleic Acids Res 50, D596–D602, doi:10.1093/nar/gkab1020 (2022).

20 Korsunsky, I. et al. Fast, sensitive and accurate integration of single-cell data with Harmony. Nat Methods 16, 1289–1296, doi:10.1038/s41592-019-0619-0 (2019).

21 Guo, C. et al. Single-cell profiling of the human decidual immune microenvironment in patients with recurrent pregnancy loss. Cell Discov 7, 1, doi:10.1038/s41421-020-00236-z (2021).

22 Denisenko, E. et al. Systematic assessment of tissue dissociation and storage biases in single-cell and single-nucleus RNA-seq workflows. Genome Biol 21, 130, doi:10.1186/s13059-020-02048-6 (2020).

23 Lotfollahi, M. et al. Mapping single-cell data to reference atlases by transfer learning. Nat Biotechnol 40, 121–130, doi:10.1038/s41587-021-01001-7 (2022).

24 Bassez, A. et al. A single-cell map of intratumoral changes during anti-PD1 treatment of patients with breast cancer. Nat Med 27, 820–832, doi:10.1038/s41591-021-01323-8 (2021).

25 Sade-Feldman, M. et al. Defining T Cell States Associated with Response to Checkpoint Immunotherapy in Melanoma. Cell 175, 998–1013 e1020, doi:10.1016/j.cell.2018.10.038 (2018).

26 Traag, V. A., Waltman, L. & van Eck, N. J. From Louvain to Leiden: guaranteeing well-connected communities. Sci Rep 9, 5233, doi:10.1038/s41598-019-41695-z (2019).

27 Xu, J. et al. ZFP541 maintains the repression of pre-pachytene transcriptional programs and promotes male meiosis progression. Cell Rep 38, 110540, doi:10.1016/j.celrep.2022.110540 (2022).

28 Li, L. et al. The XRN1-regulated RNA helicase activity of YTHDC2 ensures mouse fertility independently of m(6)A recognition. Mol Cell 82, 1678–1690 e1612, doi:10.1016/j.molcel.2022.02.034 (2022).

29 Bailey, A. S. et al. The conserved RNA helicase YTHDC2 regulates the transition from proliferation to differentiation in the germline. Elife 6, doi:10.7554/eLife.26116 (2017).

30 Jain, D. et al. ketu mutant mice uncover an essential meiotic function for the ancient RNA helicase YTHDC2. Elife 7, doi:10.7554/eLife.30919 (2018).

31 Zhang, X., Gunewardena, S. & Wang, N. Nutrient restriction synergizes with retinoic acid to induce mammalian meiotic initiation in vitro. Nat Commun 12, 1758, doi:10.1038/s41467-021-22021-6 (2021).

32 Becht, E. et al. Dimensionality reduction for visualizing single-cell data using UMAP. Nat Biotechnol, doi:10.1038/nbt.4314 (2018).

33 Han, X. et al. Mapping the Mouse Cell Atlas by Microwell-Seq. Cell 172, 1091–1107 e1017, doi:10.1016/j.cell.2018.02.001 (2018).

34 Swamy, V. S., Fufa, T. D., Hufnagel, R. B. & McGaughey, D. M. Building the mega single-cell transcriptome ocular meta-atlas. Gigascience 10, doi:10.1093/gigascience/giab061 (2021).

35 Steuernagel, L. et al. HypoMap-a unified single-cell gene expression atlas of the murine hypothalamus. Nat Metab 4, 1402–1419, doi:10.1038/s42255-022-00657-y (2022).

36 Grieshaber-Bouyer, R. et al. The neutrotime transcriptional signature defines a single continuum of neutrophils across biological compartments. Nat Commun 12, 2856, doi:10.1038/s41467-021-22973-9 (2021).

